# First identification of camel prion disease in Tataouine, Tunisia: an emerging animal prion disease in North Africa

**DOI:** 10.1101/2025.09.17.675824

**Authors:** Abdelkader Amara, Michele Angelo Di Bari, Kéfia Elmehatli, Rosalia Bruno, Rihab Andolsi, Barbara Chiappini, Ilaria Vanni, Elena Esposito, Geraldina Riccardi, Obaid Allah Ben Abid, Stefano Marcon, Atef Malek, Boubaker Ben Smida, Haykel Kessa, Walid Chandoul, Mariem Handous, Roukaya Khorchani, Romolo Nonno, Malek Zrelli, Umberto Agrimi, Gabriele Vaccari, Laura Pirisinu

**Author notes:** Correspondence should be addressed to Gabriele Vaccari or Laura Pirisinu. These authors contributed equally (Abdelkader Amara, Michele Angelo Di Bari).

## Abstract

Prion diseases are fatal neurodegenerative disorders affecting humans and animals. Among these, camel prion disease (CPrD) was recently identified in Algeria as a novel disease. In this study, we report six CPrD cases in dromedary camels (*Camelus dromedarius*) from Tunisia, providing further evidence of its occurrence in North Africa. Affected animals exhibited neurological signs and showed PrP^Sc^ accumulation in both brain and lymphoid tissues. Molecular and pathological analyses revealed features consistent with Algerian CPrD cases and distinct from classical scrapie and bovine spongiform encephalopathy. The detection of PrP^Sc^ in lymphoid organs, together with the relatively young age of some affected individuals, suggests the possibility of a contagious etiology, including potential vertical or early-life transmission mechanisms, as observed in scrapie and chronic wasting disease affecting small ruminants and cervids, respectively. These findings underscore the need for continued surveillance and further investigation into the epidemiology, transmission mechanisms and potential public health implications of CPrD.

## Introduction

Prion diseases, or Transmissible Spongiform Encephalopathies (TSEs), are invariably fatal and transmissible neurodegenerative disorders affecting both humans and animals. They are characterized by the accumulation in the central nervous system of an abnormally folded isoform (PrP^Sc^) of the host-encoded cellular prion protein (PrP^C^). According to the protein-only hypothesis, PrP^Sc^ constitutes the primary, if not exclusive, component of the infectious agent (1), and distinct conformations of PrP^Sc^ underlie the existence of different prion strains. They are associated with heritable phenotypic properties, including specific clinical signs, neuropathological patterns and biochemical characteristics (2, 3). Although prion diseases can have different origin (i.e. genetic, spontaneous/idiopathic or acquired) and occurrence (i.e. sporadic, endemic or epidemic), all forms share the fundamental feature of PrP^Sc^ misfolding and accumulation and have been historically defined based on their experimental transmissibility.

In humans, most of TSEs have idiopathic (sporadic Creutzfeldt-Jakob disease, Variably Protease Sensitive Prionopathy) or genetic origin (Fatal familial insomnia, Gerstmann-Sträussler-Scheinker disease, genetic Creutzfeldt-Jakob disease). However, those that have raised the most serious public concerns are the acquired forms, with zoonotic (variant Creutzfeldt-Jakob disease, vCJD) or iatrogenic origin (iatrogenic Creutzfeldt-Jakob disease) (4).

In animals, prion diseases are primarily acquired, such as classical scrapie in small ruminants, bovine spongiform encephalopathy (BSE) in cattle, and chronic wasting disease (CWD) in cervids (5). Nonetheless, atypical and supposed spontaneous forms, such as Nor98/atypical scrapie in small ruminants and H- and L-type BSE in cattle, have increasingly been identified, lacking clear epidemiological evidence of transmissibility under natural conditions (6–8).

Despite their rarity, prion diseases have caused significant epidemics with serious implications. Kuru, transmitted through ritual cannibalism in Papua New Guinea, was the first documented human prion epidemic (9). Later, the BSE crisis in the United Kingdom and the subsequent emergence of vCJD in humans raised major public health concerns. Molecular and biological strain typing ultimately confirmed that the prion strain causing vCJD was identical to that of BSE, thereby providing definitive evidence for a causal link and cross-species transmission (10). While control measures such as feed bans and surveillance programs mitigated the BSE epidemic, the long incubation period limited the timeliness of these interventions.

The animal TSE outbreaks, such as BSE in Europe and CWD in North America, have underscored the persistent risks associated with animal prion diseases, including their economic, ecological and zoonotic consequences. The phenomenon of cross-species transmission remains a central concern, given the permeability of species barriers. Factors such as strain properties, host compatibility and exposure dose critically influence the outcome of such events.

In recent years, novel animal prion diseases have been identified, including CWD in European cervids (11) and a prion disease, termed camel prion disease (CPrD), affecting a previously unreported host species, the dromedary camel (*Camelus dromedarius*), in Algeria (12).

While the prevalence, host range, and zoonotic potential of these emerging prion diseases remain largely unknown, they highlight the need for expanded surveillance, particularly in under-monitored regions such as Africa, where information regarding the presence and spread of animal prion diseases is limited.

We recently reported the first documented case of classical scrapie in sheep in Tunisia (13), adding to previous evidence of classical scrapie in North Africa (14).

Following the identification of CPrD in Algeria, an epidemiological surveillance network was set up in Tunisia to monitor neurological diseases in dromedary camels. The present study shows the results of investigations on suspected cases to date and reports the detection of six CPrD cases in Tunisia.

## Materials and Methods

### Animals and tissues

Over a period of approximately three years, eight dromedary camels displaying neurological signs compatible with the clinical presentation of CPrD, resulting negative for rabies, were reported (Table 1 and Suppl. Information 1). Some cases had been reported by the local breeders under the term “Medhbouba”, denoting a neurological disorder in camels. The condition is initially characterized by disorientation of the animal, which ceases to follow the flock, and subsequently accompanied by neurological signs such as head swaying movements, hyperexcitability, teeth grinding and occasionally ataxia. All animals originated from the Tataouine governorate, except for one case from the Sousse governorate and one of Algerian origin but grazing in Tunisia and collected in the Tataouine governorate (Table 1). Both frozen and formalin-fixed brain tissues were collected from these animals (Suppl. Table 1), although many samples showed evidence of tissue degradation related to storage and transportation conditions. Additionally, formalin-fixed lymph nodes were sampled from some animals: retropharyngeal lymph node (P81/9), mandibular lymph node (P81/17, P81/65), prescapular lymph node (P81/17), and unspecified lymph node from the head (P81/16).

**Table 1.**
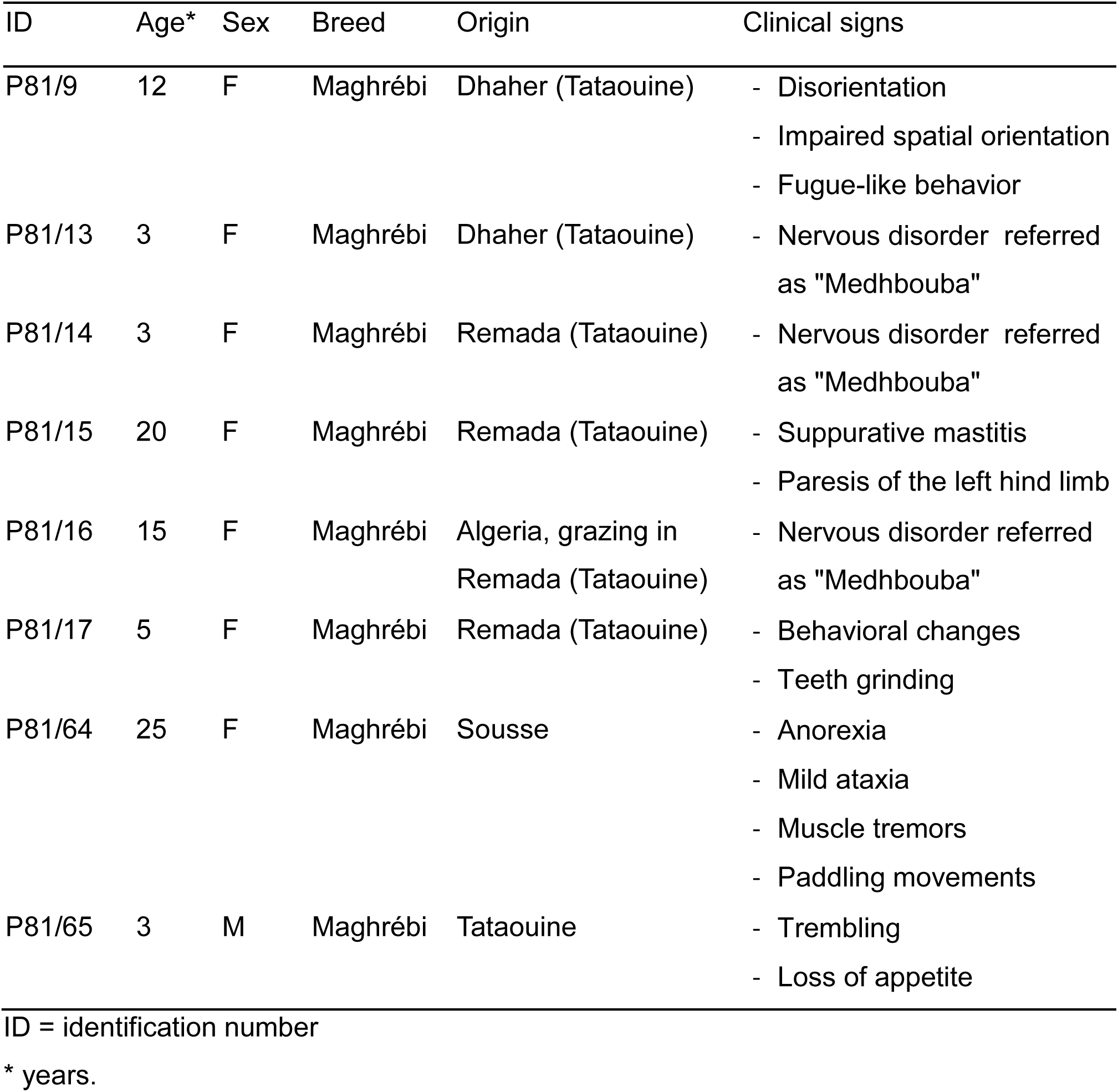
Suspect CPrD cases investigated in the study.

Brain samples were screened for the presence of rabies virus using the Fluorescent Antibody Test (FAT) and the Rabies Tissue Culture Infection Test (RTCIT) at the Rabies Laboratory of the Pasteur Institute of Tunis.

### Anti-Prion protein monoclonal antibodies

Several monoclonal antibodies (mAbs) with different epitopes were used for Western blot (WB) and immunohistochemistry (IHC): EP1802Y by Abcam (Cambridge CB2 0AX UK), SAF70, SAF84, SAF32, Sha31 were obtained by Bertin Pharma (78180 Montigny-le-Bretonneux, France), L42 and P4 by R-Biopharm (64297 Darmstadt, Germany), mAb132 by Creative Biolabs (NY 11967, USA), 9A2 and 12B2 by Wageningen Bioveterinary Research-WBWR (8221 RA Lelystad, The Netherlands).

### PrP^res^ detection

The brain tissues were analysed by a commercially available Western blot test (TeSeE Western blot; Bio-Rad Laboratories, Inc., Hercules, CA, USA) and in-house western blot protocol for detection of the protease-resistant core of PrP^Sc^ (PrP^res^). The western blot by TeSeE kit was performed as recommended by the manufacturer (Bio-Rad), using the monoclonal antibody Sha31 (AbI kit reagent). The in-house western blot diagnosis was made by a modified ISS discriminatory WB method (see below) that employed 50 µg/mL of Proteinase K (PK) and mAbs L42 and/or 12B2.

### Genotyping of dromedary camel PRNP

DNA was extracted from 100 mg of frozen brain tissue with DNeasy Blood and Tissue Kit (QIAGEN, Hilden, Germany) following the manufacturer’s instructions. The PrP gene (PRNP) coding sequence was amplified in a 50 µL final volume using 5 µL of extracted DNA, 1× AmpliTaq Gold 360 PCR Buffer (Applied Biosystems, Foster City, CA, USA), 2.5 mmol/L MgCl_2_, 1× 360 GC Enhancer, 200 µmol/L dNTPs, 0.25 µmol/L of forward (5′-GCTGACACCCTCTTTATTTTGCAG-3′) and reverse (5′-GATTAAGAAGATAATGAAAACAGGAAG-3′) primers (15), and 0.5 µL of AmpliTaq Gold 360 (Applied Biosystems), using the following amplification protocol: 5 min at 96° C; 30s at 96° C, 15s at 57° C, 9s at 72° C for 40 cycles and 4 min at 72°C. Amplicons were purified with the Illustra ExoProStar 1-Step clean-up kit (GE Healthcare Life Sciences, Little Chalfont, UK). Sequencing reactions were obtained using the BigDye Terminator v1.1 Cycle Sequencing Kit, purified using BigDye XTerminator Purification Kit, and detected with the ABI PRISM 3130 apparatus (Applied Biosystems). Sequences were analysed with SeqScape version 4 (Applied Biosystems) and compared to the wild-type allele (GenBank accession no. MF990558).

For one sample (P81/64) frozen tissue arrived at the laboratory thawed and with advanced autolytic changes. The brain material was nonetheless processed for analysis, but PrP genotype could not be determined.

### Neuropathologic and immunohistochemical analyses

Formalin-fixed brain and lymphoid samples were decontaminated with formic acid for 1 hour and then embedded in paraffin wax. Paraffin-embedded tissue blocks were cut at 5 µm for haematoxylin and eosin (H&E) staining and immunohistochemistry (IHC). IHC was performed as described previously (12), using the mAb L42. Each run included positive and negative control sections.

### Biochemical characterization of PrP^res^

PrP^res^ characterization of the positive samples was performed using the ISS discriminatory Western blot method, validated for the surveillance of animal TSEs in Europe (16). This method employs high concentrations of proteinase K for digestion, in contrast to the diagnostic WB previously applied in this study, which used 50 µg/ml PK. The protocol enables molecular typing of the protease-resistant PrP^res^ core through the combined use of monoclonal antibodies recognizing distinct epitopes across the PrP protein. Brain homogenates at 10% (wt/vol) in 100 mmol/L Tris-HCl (pH 7.4) 2% sarkosyl were incubated for 1 h at 37°C with PK (Sigma-Aldrich, St. Louis, Missouri, USA) to a final concentration of 200 µg/mL. Protease treatment was stopped with 6 mmol/L PMSF (Sigma-Aldrich). Aliquots of samples were added with an equal volume of isopropanol/butanol (1:1 vol/vol) and centrifuged at 20,000 × g for 10 min. The pellets were resuspended in denaturing sample buffer (NuPAGE LDS Sample Buffer; Life Technologies) and heated for 10 min at 95°C. We loaded each sample onto 12% bis-Tris polyacrylamide gels (Invitrogen) for electrophoresis with subsequent WB on polyvinylidene fluoride membranes using the Trans-Blot Turbo Transfer System (Bio-Rad) according to the manufacturer’s instructions. The blots were processed with several anti-PrP mAbs by using the SNAP i.d. 2.0 system (Millipore, Burlington, MA, USA) according to the manufacturer’s instructions. After incubation with the secondary antibody horseradish peroxidase-conjugated (HRP) anti-mouse immunoglobulin (Pierce Biotechnology, Waltham, MA, USA) at 1:20,000 (or goat anti-rabbit IgG (H+L) HRP at 1:10000, G21234 Thermofisher Scientific, when we used EP1802Y as primary antibody), the PrP bands were detected by using enhanced chemiluminescent substrate (SuperSignal Femto; Pierce Biotechnology) and ChemiDoc imaging system (Bio-Rad). The chemiluminescence signal was quantified by using Image Lab 6.1.0 (Bio-Rad).

## Results

### Collected samples and diagnostic evaluation

Following the first identification of a prion disease in dromedary camels in Algeria (12), investigations were conducted on the dromedary population in Tunisia. Tunisia hosts approximately 57,000 dromedary camels, about 75% of which are in the southern regions (source Direction Générale des Services Vétérinaires). For this reason, the initial investigation focused on a southern governorate (Tataouine), the southernmost region of Tunisia that borders both Libya and Algeria. Over approximately three years, eight dromedary camels were identified that exhibited neurological symptoms, behavioural changes and clinical signs consistent with previously described features of CPrD (12), raising clinical suspicion of prion disease (Table 1 and Suppl. Information 1). All animals originated from the Tataouine region, except for one case reported in the Sousse governorate and another collected in the Tataouine governorate but originating from Algeria (Table 1). The animals ranged from 3 to 25 years of age and included seven females and one male.

Brain samples from all animals underwent neuropathological examination to assess spongiform changes and were analysed for the presence of PrP^Sc^ using Western blot (WB) and immunohistochemistry (IHC). Lymph nodes were examined by IHC (Table 2).

**Table 2.**
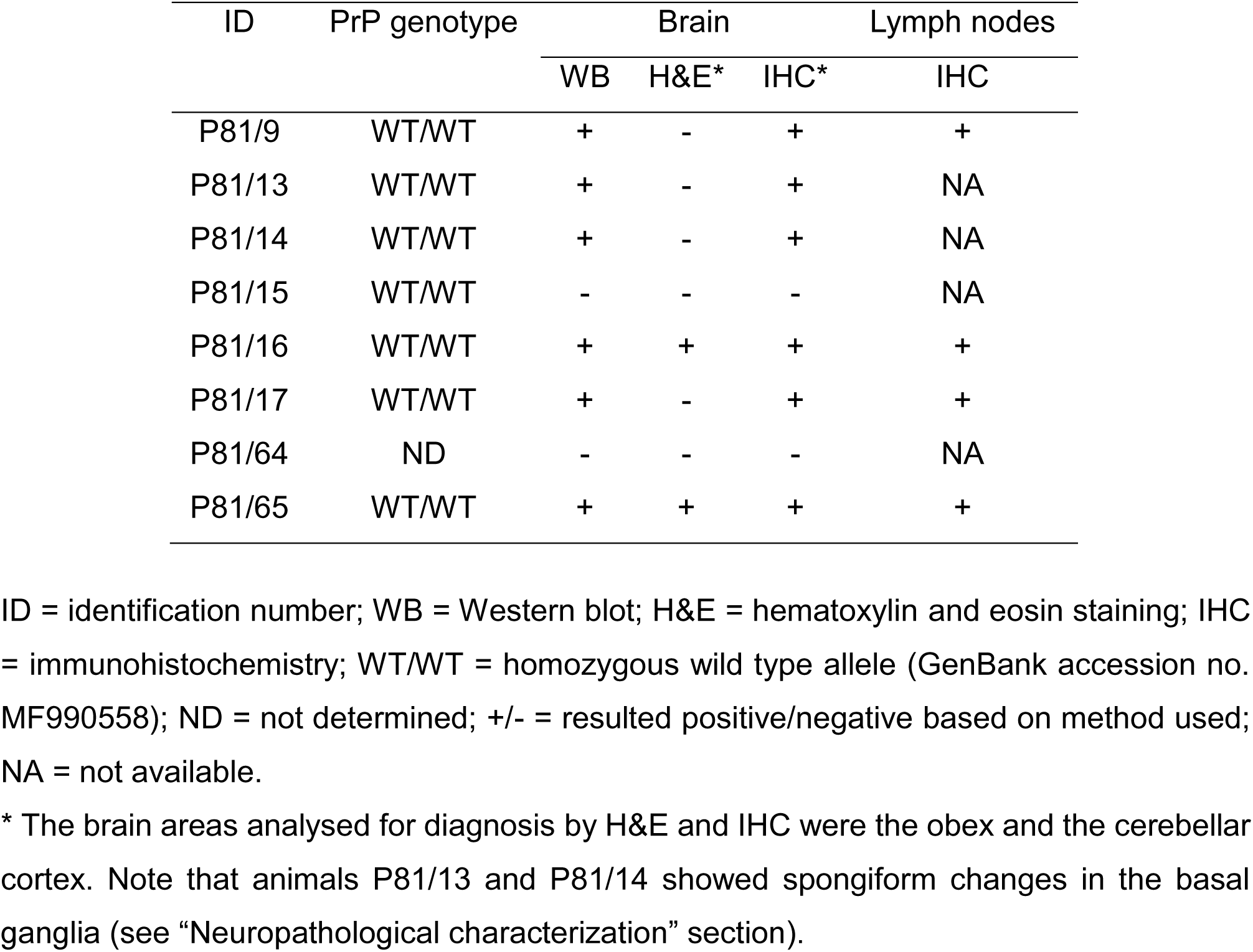
Analyses of dromedary camel samples from Tunisia.

Brain tissue from all animals was initially analyzed for PrP^Sc^ detection using a commercially available WB kit validated for small ruminants, bovines, and cervids (Figure 1A), as well as by the ISS western WB, a validated method for prion strain discrimination within the framework of the European TSE surveillance programme, with minor modifications for preliminary diagnosis. Six animals tested positive (P81/9, P81/13, P81/14, P81/16, P81/17 and P81/65) revealing the presence of the pathognomonic protease-resistant PrP^Sc^ (PrP^res^), characterized by the classical three-band pattern in the 18-30 kDa range (Figure 1, panel A). Conversely, samples P81/15 and P81/64 resulted negative. Abundant PrP^Sc^ was detected in all available brain regions of positive animals (Figure 1, panel B), providing evidence of a large involvement of several brain areas.

**Figure 1.**
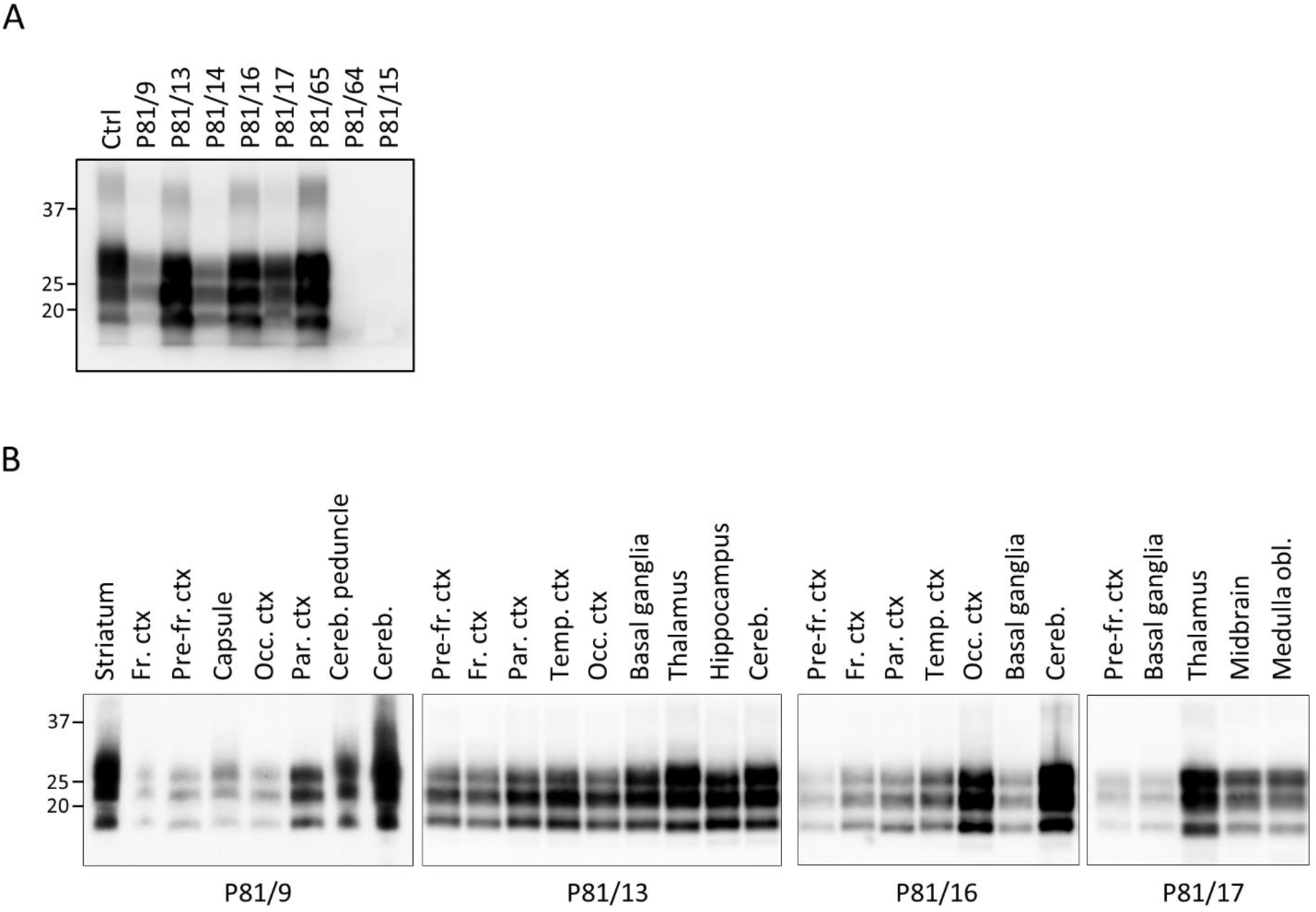
Western blot analysis of brain tissues from Tunisian dromedary camels for the detection of the proteinase K-resistant core (PrP^res^) of PrP^Sc^. A) PrP^res^ was detected using the TeSeE^TM^ Western Blot detection kit (Bio-Rad) according to the manufacturer’s instructions. The membrane was probed with the Sha31 monoclonal antibody. Loading order (left to right): classical scrapie control (Ctrl), Tunisian CPrD-positive cases (P81/9–65), and Tunisian CPrD-negative samples (P81/64 and P81/15). All samples, except for P81/64 and P81/15, were diluted 1:4 prior to electrophoresis and loaded at 3.75 mg tissue equivalent per lane. The negative-CPrD samples were loaded at 15 mg tissue equivalent per lane. Approximate molecular weights (expressed in kDa) are reported on the left of the blot. B) Representative Western blot (WB) analysis of different available brain regions from selected positive cases. Brain tissues were analyzed using the ISS WB protocol. Case identifiers are indicated below each blot, while the corresponding brain regions are labeled at the top. Abbreviations: Fr. ctx (frontal cortex), Pre-fr. ctx (prefrontal cortex), Occ. ctx (occipital cortex), Par. ctx (parietal cortex), Cereb. (cerebellum), Temp. ctx (temporal cortex). All membranes were probed with the 12B2 antibody, except for P81/17, which was probed with L42. Tissue equivalents loaded per lane: 2 mg for P81/9; 0.5 mg for P81/13, P81/16, and P81/17. Molecular weights (in kDa) are indicated on the left.

From all animals, the medulla oblongata and cerebellar cortex, target areas for animal TSE surveillance, were analysed by histopathology and IHC (Table 2). Histopathological analysis revealed spongiform changes only in animals P81/65 and P81/16, in both the medulla oblongata and cerebellar cortex, whereas spongiform change was absent in the other animals. The medulla oblongata of animal P81/65 showed intense vacuolation of the neuropil and neurons, mainly in the dorsal vagal nucleus and the nucleus of the solitary tract (Figure 2, panel A). In the obex of P81/16, rare vacuoles were found in the neuropil of the nucleus of the solitary tract. Similarly, in the molecular layer of the cerebellar cortex, intense spongiform change was observed in P81/65 (Figure 2, panel C), while only scattered vacuoles were seen in sample P81/16.

**Figure 2.**
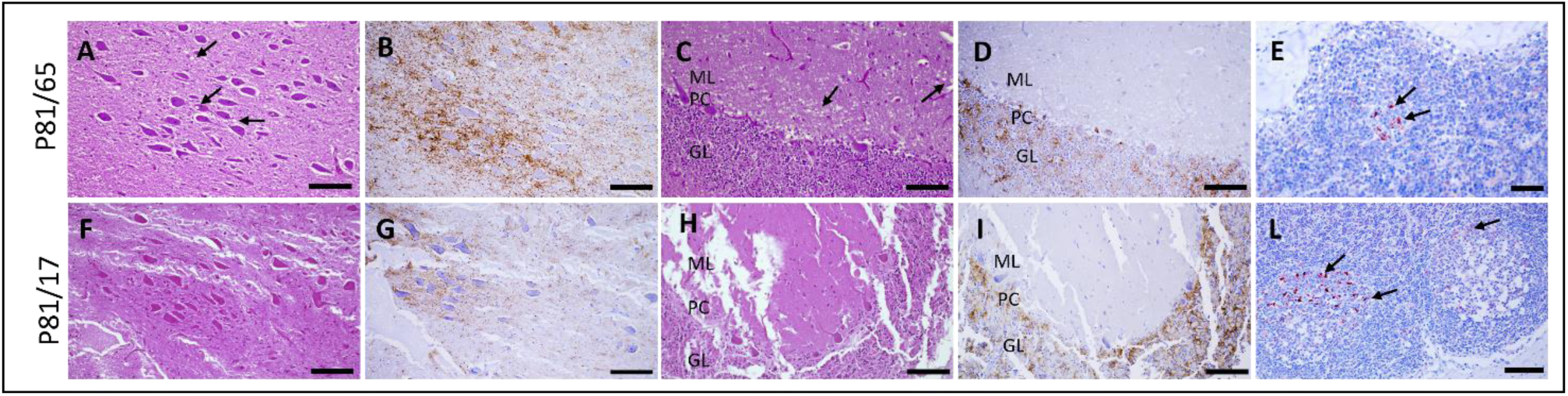
Histopathological and immunohistochemical analyses. The images show the analyses performed on the obex (A-B, F-G), cerebellar cortex (C-D, H-I) and lymph nodes (E, L) of two representative animals, P81/65 (A-E) and P81/17 (F-L). Histopathological analysis by H&E staining of the obex (A and F) and cerebellar cortex (C and H) revealed spongiform changes in the nucleus of the solitary tract (A, arrows) and in the molecular layer of cerebellar cortex (C, arrows) of P81/65, but no such alterations were observed in P81/17 (F and H). Conversely, immunohistochemistry revealed PrP^Sc^ deposition in both animals (B, D, G, I), affecting the nucleus of the solitary tract (B and G) and the cerebellar cortex (D and I). Brain sections from P81/17 showed a partial loss of tissue integrity. Immunostaining (arrows) was observed in the germinal center of both the mandibular lymph node of P81/65 (E) and the prescapular lymph node of P81/17 (L). Abbreviations: ML, molecular layer; PC, Purkinje cells; GL, granular layer. Scale bar, 50 μm in A-D and F-I; 20 μm in E and L.

Immunohistochemistry performed on the medulla oblongata and cerebellar cortex demonstrated the presence of PrP^Sc^ in all samples except P81/15 and P81/64. Notably, four animals (P81/9, P81/13, P81/14, P81/17) that did not exhibit vacuolation resulted positive by IHC, showing PrP^Sc^ deposits even in the absence of spongiform changes (Table 2 and Figure 2, panels F-I). In the obex, intense PrP^Sc^ immunoreactivity was observed in the dorsal vagal nucleus, in the nucleus of the solitary tract (Figure 2, panels B, G), as well as in the hypoglossal and olivary nuclei, and the reticular formation. The distribution of PrP^Sc^ immunostaining in the obex was similar across all cases. The cerebellar cortex displayed extensive PrP^Sc^ immunolabeling, involving the molecular and granular layers, the Purkinje cells, and the white matter (Figure 2, panels D, I). Notably, the molecular layer displayed variable intensity of PrP^Sc^ deposition across samples, with stronger immunolabeling observed in P18/13 and P81/16 compared to P81/9, P81/14, P81/17, and P81/65.

Lymph nodes were available from 4 dromedaries, all of which scored positive in the brain. Accordingly, all lymph nodes from these dromedaries revealed PrP^Sc^ deposits (Table 2), affecting both primary and secondary follicles. PrP^Sc^ deposition appeared as a reticular network within the lymphoid follicles, with intense granular depositions in tingible body macrophages within germinal centers (Figure 2, panels E, L).

Overall, the analyses performed on the collected samples showed concordance among the different techniques, identifying six positive cases out of eight suspects. In four positive animals, lymph nodes were also available and tested positive. However, four animals positive for PrP^Sc^ in the brain by both WB and IHC (including two with PrP^Sc^-positive lymph nodes) did not exhibit evident vacuolation in the obex or cerebellum. These findings support the utility and validity of both Western blot and immunohistochemistry for the diagnosis of CPrD.

It is worth noting that the two negative cases (P81/15 and P81/64) had been reported and sent to the slaughterhouse for conditions not strictly related to prion disease.

Animal P81/64 had been reported with suspected enterotoxemia (confirmed at necropsy), a condition which in dromedaries can present with neurological signs such as tremors, ataxia, aggression, hyperexcitability, convulsions, recumbency, opisthotonos, and death (17). These observations suggest that the clinical selection of suspect cases was appropriate and effective for CPrD diagnosis. However, animal P81/15 also presented with the left hind limb paresis, a symptom not strictly indicative of prion disease but relevant in the context of differential diagnosis.

Sequencing analysis of the entire PrP coding sequence revealed that the seven dromedary camels shared the same genetic background, being homozygous for the wild-type PrP allele (Table 2).

### Neuropathological characterization

Following the examination of the target areas for animal TSE surveillance (medulla oblongata and cerebellum), we extended the histopathological and immunohistochemical analysis to other brain regions available for each case, except for animal P81/65, for which only the medulla oblongata and cerebellum were available (Suppl. Table 1 and Suppl. Table 2). We thus analyzed the cerebral cortices (available from seven animals), basal ganglia (from four animals), and the pons and thalamus (available only from P81/9 and P81/15).

Histopathological analysis of the medulla oblongata and cerebellum revealed spongiform degeneration in only two of the six positive animals (P81/16 and P81/65) (Table 2). Examination of all available cerebral cortices (n = 7) showed mild spongiform changes exclusively in the temporal and occipital cortex of sample P81/16, where vacuoles were restricted to the neuropil of layers I, V, and VI (Figure 3, panel A). No spongiform changes were observed in the remaining cortical samples. In the basal ganglia (n = 4), scattered vacuoles were observed in P81/13 and P81/14, involving both white and gray matter. Spongiform change was not detected in the pons or thalamus of P81/9 and P81/15. Immunohistochemical analysis revealed no evidence of PrP^Sc^ in any of the available brain areas from samples P81/15 and P81/64, whereas all available brain regions from the other six cases resulted positive for PrP^Sc^, although cortical areas were less affected than subcortical regions. In the cortex, PrP^Sc^ deposition was predominantly localized to layers I, V, and VI (Figure 3, panels B-C). In contrast, the basal ganglia and thalamus (Figure 3, panel D) showed marked PrP^Sc^ accumulation in both gray and white matter. Multiple PrP^Sc^ deposition patterns were identified, including punctate/diffuse, intraneuronal, perineuronal, stellate, and perivascular types, affecting the neuropil, neurons, and glial cells (Figure 3, panels E–I). Notably, in the medulla oblongata, we observed a dense intra-astrocytic PrP^Sc^ deposition that filled the entire cytoplasm (Figure 3, panel L), consistent with previous reports (12).

**Figure 3.**
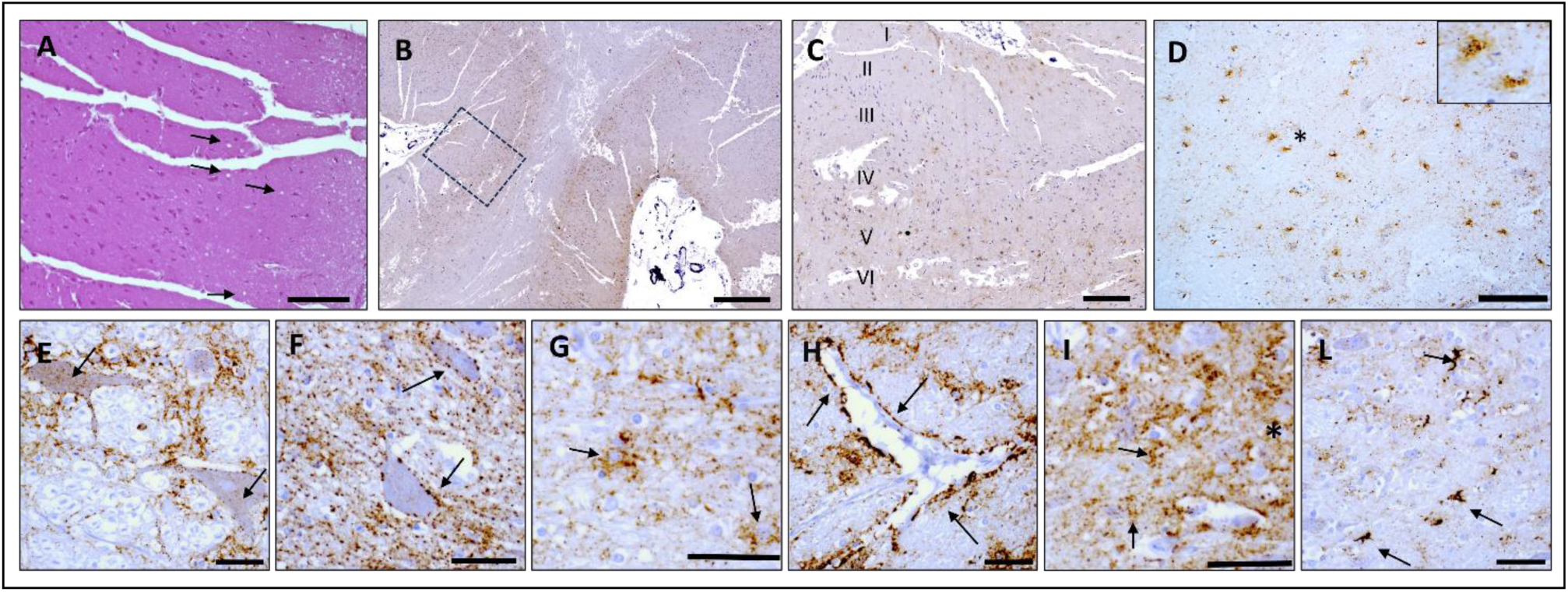
Hematoxylin and eosin staining and immunohistochemistry of brains of CPrD-affected dromedary camels. Spongiform changes in the neuropil of the occipital cortex (A) of sample P81/16. PrP^Sc^ immunostaining involving the I and V-VI layers of the prefrontal cortex (B) of sample P81/13. A magnification of the prefrontal cortex (marked by a dashed box in Panel B) is shown in the picture C. Intraglial (inset) and intraneuronal PrP^Sc^ depositions were observed in the thalamus (D) of sample P81/9. PrP^Sc^ deposition patterns observed included: intraneuronal (E, arrows), perineuronal (F, arrows), glial associated (G, arrows), perivascular (H, arrows), punctate (arrows) and diffuse (asterisk) in the neuropil (I). In the medulla oblongata, the atypical intra-astrocytic PrP^Sc^ deposition was found (L, arrows). Scale bars: 50 µm in A and D; 100 µm in B, 20 µm in C and E-L.

In conclusion, the extended histopathological analysis revealed that two animals lacking spongiform changes in the medulla oblongata and cerebellum (P81/13 and P81/14) exhibited scattered vacuoles in the basal ganglia, but not in the corresponding cortices. Immunohistochemistry across all available brain areas confirmed the diagnosis established from the medulla and cerebellum, identifying the same six positive animals out of eight examined.

### PrP^res^ characterization

Following confirmation of PrP^Sc^ presence, brain homogenates from the six positive cases underwent in-depth analysis to investigate PrP^res^ features, including the protease cleavage site, the presence of additional PrP^res^ fragments and the glycosylation patterns. A preliminary assessment of dromedary PrP^res^ reactivity with a panel of monoclonal antibodies was performed to identify suitable diagnostic tools for CPrD and to characterize PrP^res^ (Suppl. Figure 1 and Table 3).

**Table 3.**
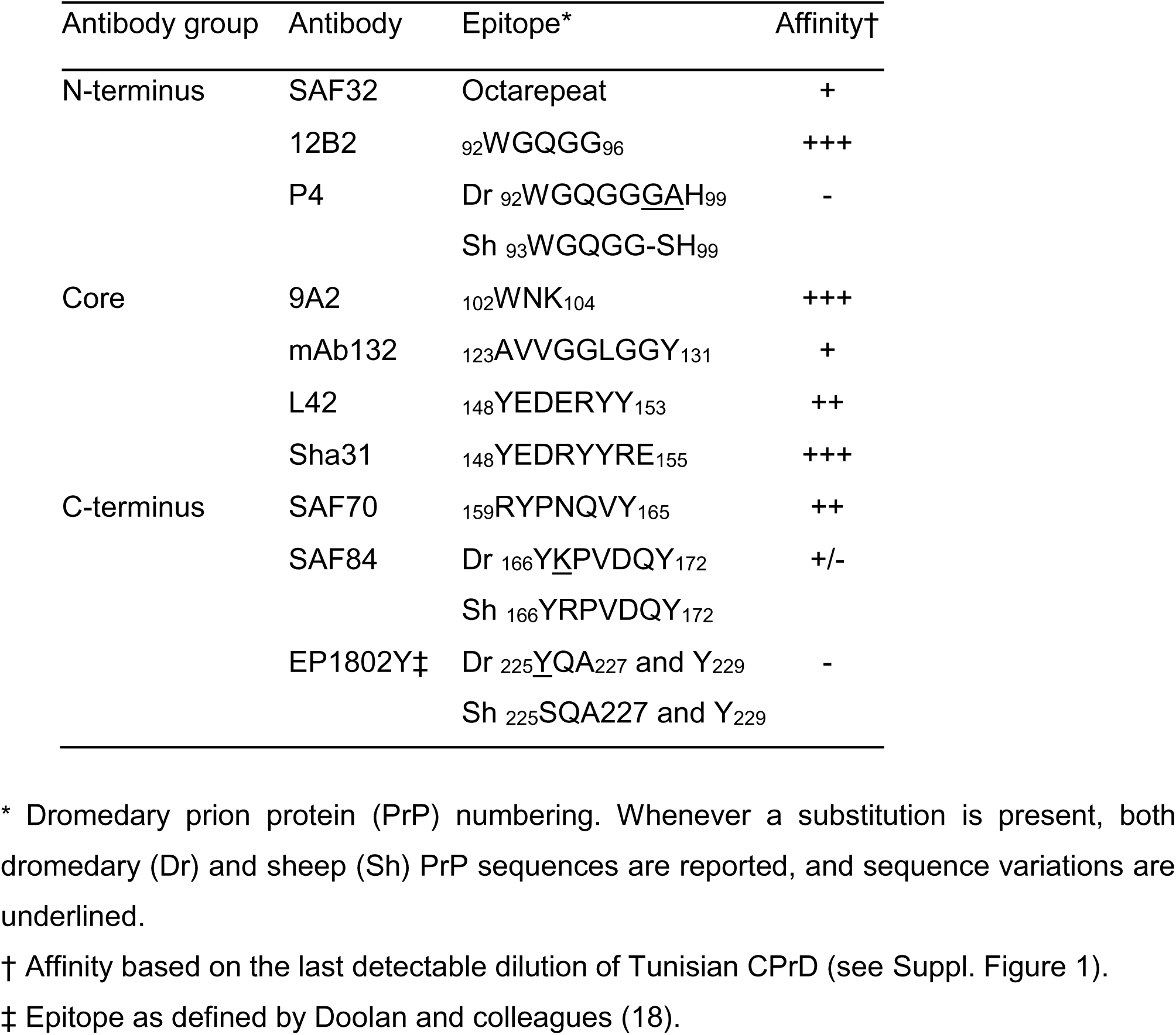
Affinity of monoclonal antibodies for dromedary camel PrP^Sc^.

To enable a detailed characterization of PrP^res^, samples were treated with high concentrations of proteinase K (PK) to clearly define the cleavage site and probed with a panel of monoclonal antibodies spanning the PrP sequence. Within each antibody group (Table 3), the antibody with the best sensitivity toward dromedary PrP^Sc^ (Suppl. Figure 1) was chosen for epitope mapping (Figure 4, panel A). Additionally, the SAF32 antibody, which targets the octarepeat region, was included to allow comparison with classical scrapie, in which this epitope is partially lost. Notably, all antibodies that showed weak or no detection of dromedary PrP target epitopes that correspond to regions with sequence variation in the dromedary PrP protein (Table 3).

**Figure 4.**
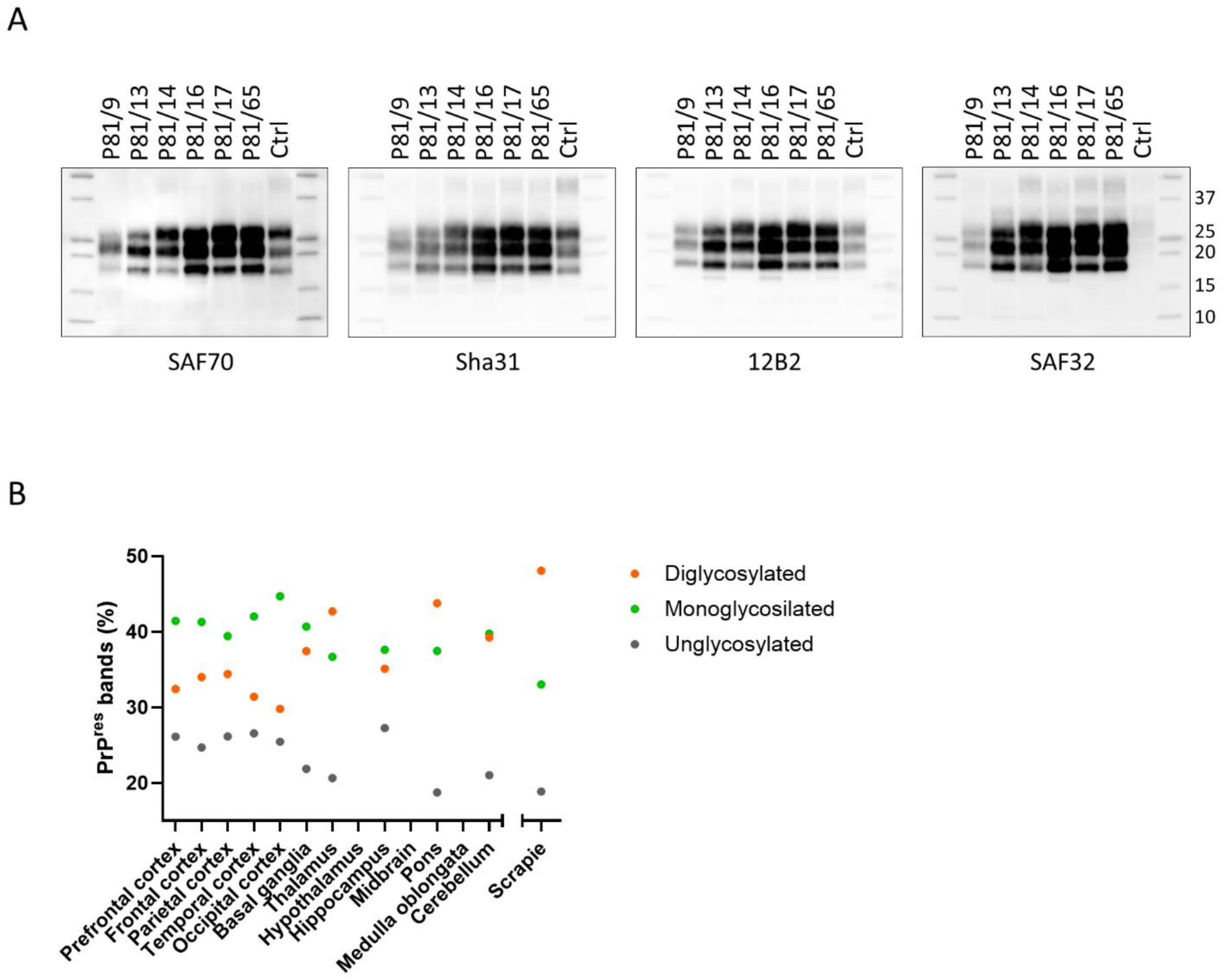
Characterization of PrP^res^ from brains of CPrD-affected dromedary camels. A) Representative replica blots showing the epitope mapping analysis of PrP^res^ from brain homogenates of positive dromedary camel cases. A classical scrapie-affected sheep sample was included as control (Ctrl) in the last lane of each blot. The brain areas analyzed for each sample were as follows: P81/9 = prefrontal cortex; P81/13 = frontal cortex; P81/14 = thalamus; P81/16 = parietal cortex; P81/17 = prefrontal cortex; P81/65 = not identifiable; scrapie = medulla oblongata. Protein standards were loaded in the first and last lanes of each blot. Membranes were probed with monoclonal antibodies indicated below each blot. B) The graph shows the relative proportion of di-, mono-, and unglycosylated PrP^res^ bands in each available brain region of animal P81/13. Scrapie is shown on the right for comparison. Quantifications were performed on membrane probed with 12B2 mAb.

PrP^res^ from dromedary isolates displayed a higher apparent molecular weight than classical scrapie control due to the preservation of the SAF32 N-terminal epitope, which is at least partially cleaved in classical scrapie (Figure 4, panel A and Suppl. Figure 1). No additional C-terminal or internal fragments were detected, as shown by the absence of signal in membranes probed with SAF70 and 12B2, respectively.

Regarding the glycosylation status, PrP^res^ from these cases exhibited lower overall glycosylation levels compared to classical scrapie (Figure 4, panel B and Suppl. Figure 2). Analysis of all available brain regions from each animal revealed minor variability; however, a consistent pattern was observed across all positive cases, with reduced PrP^res^ glycosylation in cortical areas relative to subcortical regions. Notably, the monoglycosylated isoform predominated over the diglycosylated form in the cortex (Figure 4, panel B and Suppl. Figure 2).

The PrP^res^ characteristics of these cases, electrophoretic profile, molecular weight and glycoprofile, are consistent with those previously reported in Algerian CPrD cases (12).

## Discussion

In 2018, a novel prion disease was described in dromedary camels in Algeria, based on a limited number of cases. Despite the small sample size, retrospective analyses in that study suggested that the actual prevalence might be underestimated (12). Here, we report the detection of CPrD in dromedary camels from Tunisia, providing further evidence for the geographical distribution of the disease and supporting the hypothesis of its broader presence in North Africa. Of eight clinically suspect cases investigated, six resulted positive for PrP^Sc^ in the brain, reinforcing the suspicion that CPrD is not an exceptional condition.

The molecular features of the Tunisian CPrD cases were remarkably similar to those previously described in Algerian dromedaries, and clearly distinct from classical scrapie in small ruminants and BSE in cattle. The biochemical profile of PrP^res^, including the predominance of the monoglycosylated form and the preservation of N-terminal epitope, aligns with findings from Algerian isolates and suggests the presence of the same prion strain.

The detection of PrP^Sc^ in multiple lymphoid tissues, together with the relatively young age of several affected animals, supports the hypothesis that CPrD may be a contagious prion disease. In classical scrapie in small ruminants and CWD in cervids horizontal transmission under natural conditions is likely facilitated by the involvement of lymphoreticular tissues, which might serve as peripheral sites of replication and potential sources of environmental contamination, along with the involvement and distribution in other tissues and organs (5, 19–21). In contrast, diseases such as BSE in cattle and atypical scrapie, which lack substantial lymphoid involvement, are considered inefficiently transmissible under natural conditions. In the present study, PrP^Sc^ was detected in all available lymph nodes from the positive animals, pointing toward extraneural propagation and raising concerns about potential prion shedding and environmental persistence.

We recently reported the first confirmed case of classical scrapie in sheep in Tunisia (13), in the same geographic area where CPrD cases were identified. Although there is currently no biochemical or neuropathological evidence of similarity between sheep scrapie and CPrD, potential biological relationships cannot be excluded. Bioassay experiments will be essential to clarify whether the biological properties of the prion strains involved in CPrD and scrapie share biological properties or if environmental factors might facilitate cross-species transmission or adaptation.

Taken together, the data from Algeria and Tunisia suggest that CPrD may be endemic in certain areas. Despite the limited number of reported cases so far, the absence of active surveillance programs in these countries, unlike in Europe for scrapie and BSE, raises the possibility of a higher number of undetected cases. In addition, it is worth noting the geographic proximity among the regions where cases have been identified. The positive cases described in this study originated from the Tataouine governorate, in southern Tunisia, which shares a border with the Ouargla region in southeastern Algeria, where the first CPrD cases were reported (12). While the origin of the disease remains unknown, animal movement across the Algeria-Tunisia border is a common occurrence. Indeed, one of these cases was of Algerian origin and had been grazing in the Tataouine area. In these regions, dromedaries are frequently raised under extensive pastoral systems, in which animals range freely across desert areas, sometimes covering distances of 100-150 km in search of water and food. Under such conditions, direct or indirect contact between Tunisian and Algerian camels, as well as among animals from different regions or countries, cannot be excluded, with potential implications for disease circulation.

Despite the typically long incubation periods associated with prion diseases, often spanning several years, several positive cases were notably young. Dromedaries reach sexual maturity around 3–5 years of age, and in our study, clinical signs were observed in animals as young as 3 years old. This may be due to high levels of environmental exposure and/or exposure at an early age. Indeed, several studies suggest that extensive environmental contamination, that implies high infectious dose, can lead to a shortened incubation period in prion diseases (22, 23). These findings are also consistent with patterns observed in classical scrapie and CWD, where early-life exposure (through fluids, placenta, colostrum, and milk) along with vertical transmission, contributes to the accumulation of infected individuals at a relatively young age (20, 24–33).

The potential impact of CPrD on camel population health and production is of particular concern. Dromedaries are long-lived animals with extended reproductive and productive lifespan and are critical to the livelihoods of pastoralist communities in arid regions. A transmissible prion disease in this species could lead to significant economic losses due to premature death, culling and reduced productivity in terms of milk and meat. The identification of CPrD in regions where camels play a central role in food production and local economies highlights the need for a thorough understanding of the disease’s epidemiology, transmission mechanisms and long-term impact on animal health.

The emergence of a new prion disease in animals also raises concerns for public health. To date, only BSE has demonstrated zoonotic potential among animal prion diseases. However, the BSE crisis has established a precedent for the need to rigorously monitor all emerging TSEs in animals. The long incubation period and lack of early clinical markers complicate surveillance and risk assessment. Notably, in the BSE/vCJD context, the human epidemic peaked roughly a decade after the peak of the cattle epidemic, highlighting the latency and unpredictability of zoonotic prion diseases (5).

Our data suggest that camel prion disease should be regarded as a novel emerging infectious disease in North Africa, whose prevalence, transmission routes, biological properties and interspecies transmission potential remain largely unknown. The identification of PrP^Sc^ in lymphoid tissues, the detection of multiple cases in a single region, and the age of affected animals all point toward an infectious etiology that merits further investigation.

Importantly, this study demonstrates that standard diagnostic tools (including commercial kits, Western blot and immunohistochemistry protocols) and widely used monoclonal antibodies such as Sha31, 12B2, SAF32, and L42 are suitable for the detection and characterization of PrP^Sc^ in CPrD. Overall, this shows that reliable and accessible methodologies are available to support future efforts in surveillance and diagnosis of prion diseases in dromedary camels.

It is noteworthy that, in some cases, we observed the absence or a minimal presence of spongiform change alongside evident accumulation of PrP^Sc^, as demonstrated by both immunohistochemistry and Western blot. Several studies have shown that spongiform change and PrP^Sc^ deposition in the brain may represent independent processes, without consistent temporal or spatial association, and the relationship between PrP^Sc^ accumulation, vacuolation and neurotoxicity in prion diseases is still not fully understood (34–40). Moreover, as also observed in other animal prion diseases, such as classical scrapie and CWD (23, 41), PrP^Sc^ can be detected earlier than histopathologic vacuolation, supporting and recommending the use of immunohistochemistry as a diagnostic tool.

All animals analyzed in this study were homozygous for the wild-type PRNP allele. Given the established role of host genetics in scrapie susceptibility, future studies to characterize PRNP polymorphisms in the dromedary population may help identify alleles associated with resistance and support genetic selection as a potential control strategy. However, available data suggest that PRNP variability in dromedaries appears to be limited overall, with only two non-synonymous polymorphisms having been identified in Algerian animals (42) but none detected in Ethiopian populations (43).

Given the potential implications for animal and human health, as well as the possible socio-economic impact in regions where dromedaries play a key role, ongoing surveillance and in-depth characterization of CPrD are essential to enhance our comprehension of its epidemiology, host range and zoonotic potential.

## Acknowledgments

We would like to thank Professor Baaissa Babelhadj, who first described CPrD, for providing us with the Algerian dromedary samples. This contribution was essential, as it enabled a direct comparison with the Tunisian cases included in our study.

This study was supported by the Istituto Superiore di Sanità, Independent Research Programme (grant no. ISS20-97e1d82bda5a), and by the La Sapienza University of Rome and Istituto Superiore di Sanità, Italy-Africa PhD Programme (grant no. 5165).

