## Supplementary Information for "First identification of camel prion disease in Tataouine, Tunisia: an emerging animal prion disease in North Africa"

**Supp. Information.** Clinical Presentation and animal information

1) Animal ID P81/9. The first case involved a 12-year-old female dromedary from a herd of 72 animals raised under extensive farming conditions in the Dhaher and Ouaara grazing areas without dietary supplementation. This animal displayed disorientation, a lack of spatial awareness, and fugue-like behavior approximately 15 days before being slaughtered at a slaughterhouse in the Tataouine region.
2) Animal ID P81/13. A 3-year-old female dromedary was identified at the same slaughterhouse and a few months later of Case 1. The animal exhibited nervous disorder locally referred to as "Medhbouba", characterized by initial disorientation, loss of flock-following behaviour, head swaying, hyperexcitability, teeth grinding, and ataxia.
3) Animal ID P81/14. A 3-year-old female dromedary, also diagnosed at the same slaughterhouse and time as Case 2, displayed similar nervous disorder ("Medhbouba"), reinforcing the suspicion of CPrD.
4) Animal ID P81/15. A 20-year-old female dromedary presented with suppurative mastitis and paresis of the left hind limb.
5) Animal ID P81/16. This case involved a 15-year-old female dromedary of Algerian origin, grazing in Tunisia. The animal was referred as a "Medhbouba" case by the herder. At the ante-mortem examination, the veterinarian observed teeth grinding, head swaying, a staring gaze and failure to follow conspecifics.
6) Animal ID P81/17. Within a herd of 60 animals, a 5-year-old suckling female fed straw and concentrate for four months before being released to pasture. This animal exhibited behavioral changes, such as avoiding the herd and grinding her teeth. Based on these observations, a decision was made to cull the animal, which was subsequently slaughtered.
7) Animal ID P81/64. A 25-year-old female dromedary from the Sousse region exhibited clinical signs including a three-day history of anorexia, mild ataxia, and neurological manifestations consisting of muscle tremors and paddling movements during the final day. In addition to CPrD, a presumptive diagnosis of enterotoxemia had been made prior to death. The suspicion of enterotoxemia was confirmed at necropsy, which revealed digestive acidosis associated with hepatorenal degeneration, findings consistent with enterotoxemia.
8) Animal ID P81/65. The last case involved a 3-year-old male from the Tataouine region reported to exhibit trembling and a loss of appetite.
