## Supplementary Table 1 for "First identification of camel prion disease in Tataouine, Tunisia: an emerging animal prion disease in North Africa"

**Supp. Table 1.** Collected samples

| **Brain area** | **P81/9** | | **P81/13** | | **P81/14** | | **P81/15** | | **P81/16** | | **P81/17** | | **P81/64** | | **P81/65** | |
|---|---|---|---|---|---|---|---|---|---|---|---|---|---|---|---|---|
| | FF | F | FF | F | FF | F | FF | F | FF | F | FF | F | FF | F* | FF | F* |
| Prefrontal cortex | | X | X | X | X | | X | | | X | | X | | | | |
| Frontal cortex | | X | X | X | X | X | X | X | | X | | | | | | |
| Parietal cortex | X | X | | X | | X | | | | X | | | | | | |
| Temporal cortex | X | | | X | | X | X | | X | X | | | | | | |
| Occipital cortex | X | X | | X | | | | | X | X | X | | X | | | |
| Basal ganglia | X | X | X | X | X | X | X | | | X | | X | | | | |
| Thalamus | X | | | X | | X | X | | | | | X | | | | |
| Hypothalamus | | | | | | X | | | | | | | | | | |
| Hippocampus | | | | X | | X | | | | | | | | | | |
| Midbrain | | | | | | X | | | | | | X | | | | |
| Pons | X | | | X | | | X | | | | | | | | | |
| Medulla oblongata | X | | X | | X | | X | X | X | | X | X | X | | X | |
| Cerebellum | X | X | X | X | X | X | X | | X | X | X | | X | | X | |

FF: formalin-fixed; F: frozen

* Unidentifiable areas: tissue received thawed and exhibiting autolytic changes. The brain material was nonetheless processed for analysis.
