## Supplementary Table 2 for "First identification of camel prion disease in Tataouine, Tunisia: an emerging animal prion disease in North Africa"

**Suppl. Table 2.** Results of histopathological and immunohistochemical analyses.

| **Brain area** | **P81/9** | | **P81/13** | | **P81/14** | | **P81/15** | | **P81/16** | | **P81/17** | | **P81/64** | | **P81/65** | |
|---|---|---|---|---|---|---|---|---|---|---|---|---|---|---|---|---|
| | H&E | IHC | H&E | IHC | H&E | IHC | H&E | IHC | H&E | IHC | H&E | IHC | H&E | IHC | H&E | IHC |
| Prefrontal cortex | | | - | + | - | + | - | - | | | | | | | | |
| Frontal cortex | | | - | + | - | + | - | - | | | | | | | | |
| Parietal cortex | - | + | | | | | | | | | | | | | | |
| Temporal cortex | - | + | | | | | - | - | + | + | | | | | | |
| Occipital cortex | - | + | | | | | | | + | + | - | + | - | - | | |
| Basal ganglia | - | + | + | + | + | + | - | - | | | | | | | | |
| Thalamus | - | + | | | | | - | - | | | | | | | | |
| Pons | - | + | | | | | - | - | | | | | | | | |
| Medulla oblongata | - | + | - | + | - | + | - | - | + | + | - | + | - | - | + | + |
| Cerebellum | - | + | - | + | - | + | - | - | + | + | - | + | - | - | + | + |

H&E = hematoxylin and eosin staining; IHC = immunohistochemistry
