## Supplementary Figure 1 for "First identification of camel prion disease in Tataouine, Tunisia: an emerging animal prion disease in North Africa"

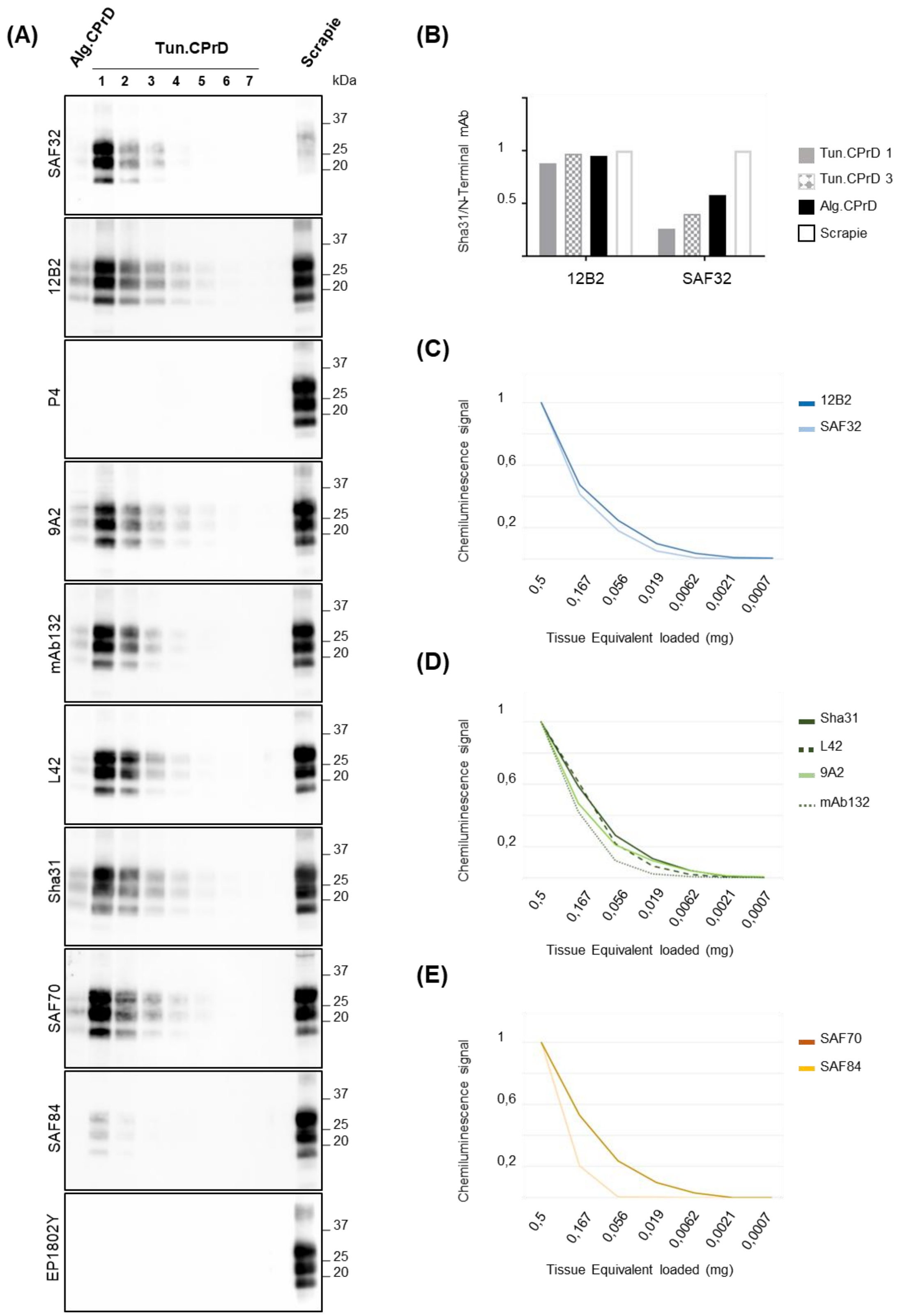


**Supp. Figure 1.** Western blot analysis of Tunisian and Algerian CPrD and evaluation of mAbs sensitivity. A) Representative western blot of PK-treated (50 µg/ml) brain homogenate of dromedary camel CPrD isolates from Algeria (Alg. CPrD) (12) and Tunisia (Tun. CPrD), and sheep classical scrapie. Replica blots were probed with different monoclonal antibodies (mAbs), as indicated on the left of each blot. The Tunisian CPrD sample was subjected to a 1:3 serial dilution (indicated as 1 to 7 in the blots). Tissue equivalents (TE) loaded per lane were 0.9 mg for Alg. CPrD, 0.5 mg for the first dilution point of the Tunisian CPrD (indicated as 1), and 0.5 mg for classical scrapie. The molecular weights (expressed in kDa) are indicated on the right of each blot. Based on the titration results, mAbs used in epitope mapping can be divided into 4 different groups based on the ability to detect $PrP^{res}$: +/-) SAF84, which detects CPrD $PrP^{res}$ only until the $2^{nd}$ dilution (0.167 mg TE loaded); +) SAF32 and mAb132 able to detect until the $3^{rd}/4^{th}$ dilution (between 0.056 mg and 0.019 mg TE loaded); ++) SAF70 and L42 which detect until the $4^{th}/5^{th}$ dilution (between 0.019 mg and 0.0062 mg TE loaded); +++) Sha31, 9A2 and 12B2 which can detect until the $5^{th}/6^{th}$ dilution (between 0.0062 mg and 0.0021 mg TE loaded). Note that the Algerian CPrD sample is characterised by an extremely low amount of $PrP^{Sc}$, comparable to the amount detected in the third dilution of the Tunisian CPrD sample. B) Graph depicting the antibody ratio (Sha31/N-Terminal Ab) of the Algerian CPrD (Alg. CPrD), undiluted Tunisian CPrD (Tun. CPrD, ID 1 in the blots in panel A), and Tunisian CPrD diluted 1:9 (Tun. CPrD, ID 3) relative to the Sha31/N-terminal Ab ratio of control scrapie. The ratio using 12B2 as N-terminal mAb is reported on the left, while using SAF32 on the right. The Sha31/12B2 ratio showed values close to 1 for all the samples, indicating the conservation of the 12B2 epitope in CPrD $PrP^{res}$. Importantly, the Sha31/SAF32 ratio was extremely low, thus indicating that the SAF32 epitope is preserved upon PK cleavage in CPrD while being partially cleaved in classical scrapie. C, D, E) Graphs depicting the chemiluminescence signal intensity obtained with antibodies used in panel A, across serial dilutions of the Tunisian CPrD sample. The y-axis represents the ratio of the chemiluminescence signal at each dilution relative to the signal at the first dilution, which is set as 1. The x-axis represents the tissue equivalents loaded (expressed in mg) in each dilution. The antibodies are grouped based on their epitope location on PrP, i.e., N-terminal region (C), core region (D), and C-terminal region (E). Among the antibodies targeting the N-terminal region of PrP (C), SAF32 shows a rapid signal decline and lower ratio values compared to 12B2 up to the $4^{th}$ dilution, where its signal tends to zero. In contrast, 12B2 still maintains a detectable signal up to the $6^{th}$ dilution. D) In line with the qualitative classification based on the last detectable dilutions (A) and similarly to mAb 12B2, antibodies 9A2 and Sha31 maintain a detectable signal at extreme high dilutions, in contrast to mab132 and L42 for which the ratios tend to zero at the $4^{th}$ and $5^{th}$ dilution, respectively. Note that, despite 9A2 shows a more rapid signal decline than L42 in the first dilutions, an inversion of the chemiluminescence ratio of the two antibodies is observed from the $4^{th}$ dilution onwards. Finally, among mAbs targeting the C-terminal region of PrP (E), SAF84 signal dramatically decreased at the second dilution point, dropping to nearly 0 at the third dilution, in contrast to SAF70 whose chemiluminescence ratio tends to zero only at the $5^{th}$ dilution.
