## Supplementary Figure 2 for "First identification of camel prion disease in Tataouine, Tunisia: an emerging animal prion disease in North Africa"

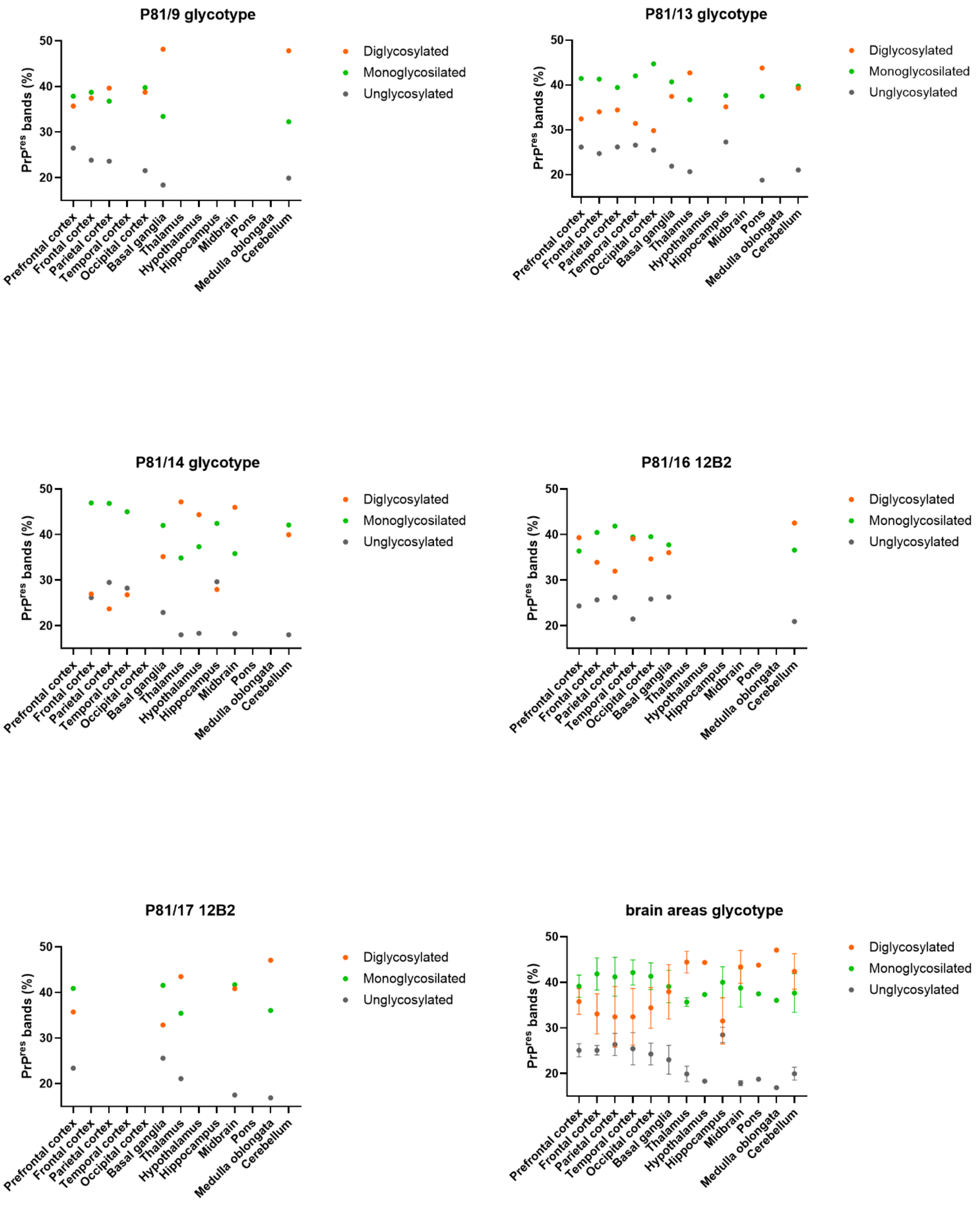


**Supp. Figure 2.** Analysis of glycotipe of $PrP^{res}$.

The graphs show the relative proportion of di-, mono-, and unglycosylated $PrP^{res}$ bands in each available brain region of all positive dromedary camels. The last graph shows the mean and standard deviation of $PrP^{res}$ forms from all animals per each brain region. Quantifications were performed on membranes probed with 12B2 mAb.
